# Fully synthetic hydrogels promote robust crypt formation in intestinal organoids

**DOI:** 10.1101/2024.07.06.602364

**Authors:** Ella A Hushka, Michael R Blatchley, Laura J Macdougall, F Max Yavitt, Bruce E Kirkpatrick, Kaustav Bera, Peter J Dempsey, Kristi S Anseth

**Affiliations:** Department of Chemical and Biological Engineering, University of Colorado Boulder, Boulder, CO 80309, USA; BioFrontiers Institute, University of Colorado Boulder, Boulder, CO 80309, USA; Medical Scientist Training Program, University of Colorado Anschutz Medical Campus, Aurora, CO 80045, USA; Section of Developmental Biology, Department of Pediatrics, University of Colorado, Denver, CO 80045, USA

## Abstract

Initial landmark studies in the design of synthetic hydrogels for intestinal organoid culture identified precise matrix requirements for differentiation, namely decompression of matrix-imposed forces and supplementation of laminin. But beyond stating the necessity of laminin, organoid-laminin interactions have gone largely unstudied, as this ubiquitous requirement of exogenous laminin hinders investigation. In this work, we exploit a fast stress relaxing, boronate ester based synthetic hydrogel for the culture of intestinal organoids, and fortuitously discover that unlike all other synthetic hydrogels to date, laminin does not need to be supplemented for crypt formation. This highly defined material provides a unique opportunity to investigate laminin-organoid interactions and how it influences crypt evolution and organoid function. Via fluorescent labeling of non-canonical amino acids, we further show that adaptable boronate ester bonds increase deposition of nascent proteins, including laminin. Collectively, these results advance the understanding of how mechanical and matricellular signaling influence intestinal organoid development.

## MAIN

Intestinal organoids hold unparalleled biomimicry and consequently great promise for studying organogenesis, modeling disease and screening drug candidates *in vitro*. Traditional intestinal organoid culture methods rely on the protein rich, biological matrix, Matrigel. Matrigel robustly promotes intestinal organoid formation and differentiation^1,2^, but because of its limited tunability and high concentration of proteins and growth factors, including extracellular matrix (ECM) proteins like laminin, the matrix limits the ability to test complex biochemical and mechanical hypotheses. In recent years there has been growing interest in utilizing synthetic materials to culture intestinal organoids to probe organoid-matrix interactions and establish culture platforms with enhanced translational potential.

Amongst the first of these efforts, Gjorevski *et al*. used synthetic, hydrolytically softening hydrogels and demonstrated that the organoids required two phenomena to support differentiation and crypt formation: (1) a decompression of forces, in this specific instance via matrix degradation, and (2) supplementation of full-length laminin-111 into the otherwise synthetic matrix.^3^ The advent of matrices with controlled degradation and dynamic matrix mechanical properties has allowed for investigation into how mechanical cues like matrix degradation influence organoid growth and differentiation.^3-6^ More recently, Chrisnandy *et al*. and Elosegui-Artola *et al*. utilized viscoelastic hydrogels to promote degradation independent crypt formation.^7,8^ However, these systems still required the addition of laminin for robust crypt formation. The former implemented a synthetic dynamic matrix with reversible triple hydrogen bonds and supplemental laminin to culture intestinal organoids, while the later utilized an alginate hydrogel supplemented with laminin rich Matrigel to achieve mature organoids. Beyond stating that organoid differentiation requires laminin, laminin has remained largely unstudied in organoid cultures despite the intriguing, ubiquitous differentiation requirement for exogenous laminin. Largely, these studies have been impeded by the laminin rich materials most used to culture intestinal organoids.

The most widely used matrix, Matrigel, is approximately 60% laminin^9^ and consequently floods organoid cultures with laminin, making it difficult to deconvolute user-imposed vs nascently deposited laminin interactions. Additionally, this high concentration of laminin introduces technical limitations for imaging, such as poor antibody penetration with antibody saturation occurring after just a few z planes.^10^ These limitations eliminate the ability to effectively image, and thus study, laminin-organoid interactions. The introduction of synthetic matrices for organoid growth has allowed some study of organoid-matrix interactions, because each component of these engineered matrices is strictly user-determined.^10^ However, the ubiquitous requirement of exogenous laminin again limits organoid-laminin interaction studies, as it is difficult to image and differentiate between user supplied versus nascent laminin. Even if this issue is overcome with advanced imaging techniques, the supplemented laminin introduces the unknown influence of exogenous laminin on organoid protein interactions and signaling. Lastly, the relatively uniform distribution of exogenous laminin throughout a hydrogel lacks physiological relevance, as laminin *in vivo* is localized to regions of the crypt-villus structure.^11,12^

Ultimately, the inability to decouple the influence of nascent and user-supplied laminin limits the study of laminin-organoid interactions. An improved understanding of organoid-matrix interactions will instruct improved material design to actualize clinical translation of intestinal organoids. To that end, we exploited a fully synthetic, fast-relaxing, boronate ester-based covalent adaptable network (CAN) and hypothesized that this rate of stress relaxation would still promote organoid differentiation without a change in network modulus. Most notably, however, we discovered this system does not require supplemental laminin for differentiation, and, to our knowledge, this system is the first fully synthetic material to achieve organoid morphogenesis and crypt formation in this manner. This phenomenon enabled our study of cell-secreted laminin organoid interactions, as compared to the exclusively studied exogenous laminin addition.

## RESULTS

### Boronate ester-based hydrogels rapidly relax matrix stress

To begin the study of organoid-matrix interactions, boronate ester-based hydrogels were synthesized using three, 20 kDa, 8-arm poly(ethylene glycol) (PEG) macromers. The first macromer is an 8-arm PEG dibenzocyclooctyne (DBCO). The second macromer has approximately six arms functionalized with nitrodopamine (ND) and two arms functionalized with azide groups. The final macromer has approximately six arms functionalized with fluorophenyl boronic acid (FPBA) and two arms functionalized with azide groups.^13^ Upon mixing, the DBCO and azide groups rapidly and permanently crosslink through a stabilizing strain promoted azide-alkyne cycloaddition (SPAAC) reaction **(Fig. 1a)**. Concurrently, the FPBA and ND groups form dynamic, covalent adaptable bonds that enable stress relaxation and viscoelasticity^13^ **(Fig. 1b)**. These reactions form a network that offers both stability and a rapid relaxation of stress without a change in network modulus (**Fig. 1c**). Boronate ester hydrogels relax 80% of the network stress on the order of seconds, regardless of macromer weight percent and modulus **(Fig. 1d,e)**. Of the dynamic covalent chemistries, the boronate ester bond type is the fastest adapting^14^ and in our system relaxes stress faster, but to the same final extent as Matrigel (data for Matrigel mined from Chrisnandy *et al*.^*8*^) (**Fig. 1d**). By varying the macromer weight percent from 3.8 to 4.6 total PEG wt%, the modulus of the matrix was varied from 500 to 2000 Pa **(Fig. 1f)**. Intestinal organoids were encapsulated as colonies in hydrogels ranging from 3.8 to 4.6 total PEG wt% and maintained in growth media for three days and then fixed. **(Fig. 1g)**. Studies with organoids were carried out using the 4.2 wt% formulation (∼1 kPa), as this condition best supported organoid viability, which was marked by the presence of epithelial polarization visualized by F-actin localization (**Fig. 1h-j**).

**Fig 1.**
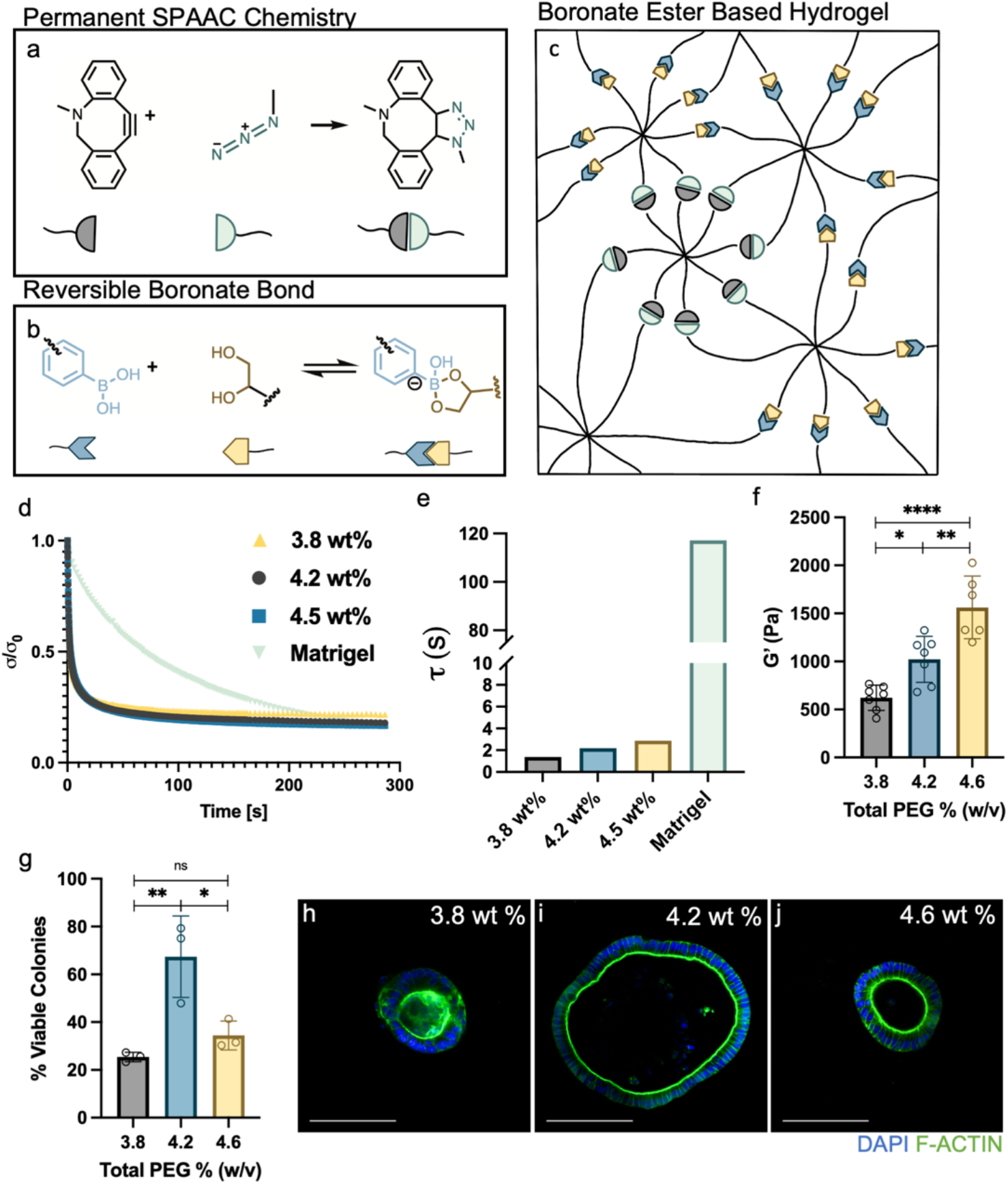
Boronate ester PEG hydrogel chemistry. **a)** Schematic of permanent strain promoted azide alkyne cycloaddition (SPAAC) reaction between a dibenzocyclooctyne (gray) and an azide group (green) **b)** a schematic of the reversible covalent adaptable bond between a fluorophenylboronic acid group (blue) and a nitrodopamine (yellow) **c)** representation of the boronate ester hydrogel network composed of an 8-arm PEG DBCO, an 8-arm PEG with 2 arms functionalized with azide groups and 6 arms functionalized with fluorophenylboronic acid, and an 8-arm PEG with 2 arms functionalized with azide groups and 6-arms functionalized with nitrodopamine. **d)** Boronate ester hydrogels relax stress rapidly, with 80% of the network stress relaxed on the order of seconds regardless of macromer weight percent. **e)** The relaxation time constant, τ, for each condition was calculated by curve fitting the averages of normalized stress data from (d) to a one component Kohlrausch-Williams-Watts function in Matlab. The hydrogels relax faster than Matrigel relax but to the same final extent (*n* = 3 for 4.2 wt%, 4.5 wt% and Matrigel, *n* = 1 for 3.8 wt %; data for Matrigel mined from Chrisnandy *et al*. **f)** Boronate ester hydrogel moduli are tuned by varying macromer weight percent (*n* = 7 for 3.8, 4.2 wt%, *n* = 6 for 4.6 wt%, mean ± s.d., \**P* ≤ 0.05, *\*\*P* ≤ 0.01, *\*\*\*\*P* ≤ 0.0001, one-way ANOVA). **g)** Intestinal organoid viability was highest in 4.2 wt% hydrogels and was used in subsequent studies (*n* = 3, ns ≥ 0.05, mean ± s.d., \**P* ≤ 0.05, *\*\*P* ≤ 0.01, one-way ANOVA) **h-j)** Viability of colonies was assessed by visualization of epithelial polarization as marked by F-actin localization (green) Scale bar 100 μm.

### Stress relaxing hydrogels promote robust crypt formation, even in the absence of supplemented laminin

To assess whether the uniquely rapid stress relaxation properties of the boronate ester-based hydrogels were sufficient to promote differentiation without a change in network modulus, intestinal organoid colonies were encapsulated in boronate ester hydrogels and immediately switched to differentiation media. After 48 hours, organoids were fixed and immunostained to evaluate for differentiated organoids, marked by the emergence of intestinal crypts containing Paneth cells that secrete lysozyme (**Fig. 2a)**. Crypt forming efficiency (CFE) was defined as the percent of colonies that formed crypts and organoids encapsulated in boronate ester-based hydrogels demonstrated robust CFEs **(Fig. 2b)**. Of great note, we find that organoids cultured in boronate ester hydrogels *without* exogenously added laminin (-Lam, **Fig. 2c**) produced differentiated organoids at a rate indistinguishable from those with laminin (+Lam, **Fig. 2d**). This is a significant result, as beyond requiring a decompression of forces, intestinal organoid differentiation in synthetic matrices (elastic or viscoelastic) have universally required laminin supplementation to promote robust crypt formation. To our knowledge, this is the first instance where the absence of laminin in the matrix does not lead to a substantial decrease or complete elimination of crypts.^3-8,15^ Intriguingly, the two publications with stress relaxing matrices used to culture intestinal organoids still required the addition of laminin for robust crypt formation.^7,8^ Notably, the dynamics of relaxation in the boronate ester hydrogels are substantially different from previously employed stress relaxing matrices and thus may explain this discrepancy. This novel finding suggests to us that the viscoelastic boronate ester system must uniquely influence matricellular signaling compared to other synthetic matrices and compels investigations into the dynamics of organoid-matrix signaling.

**Fig 2.**
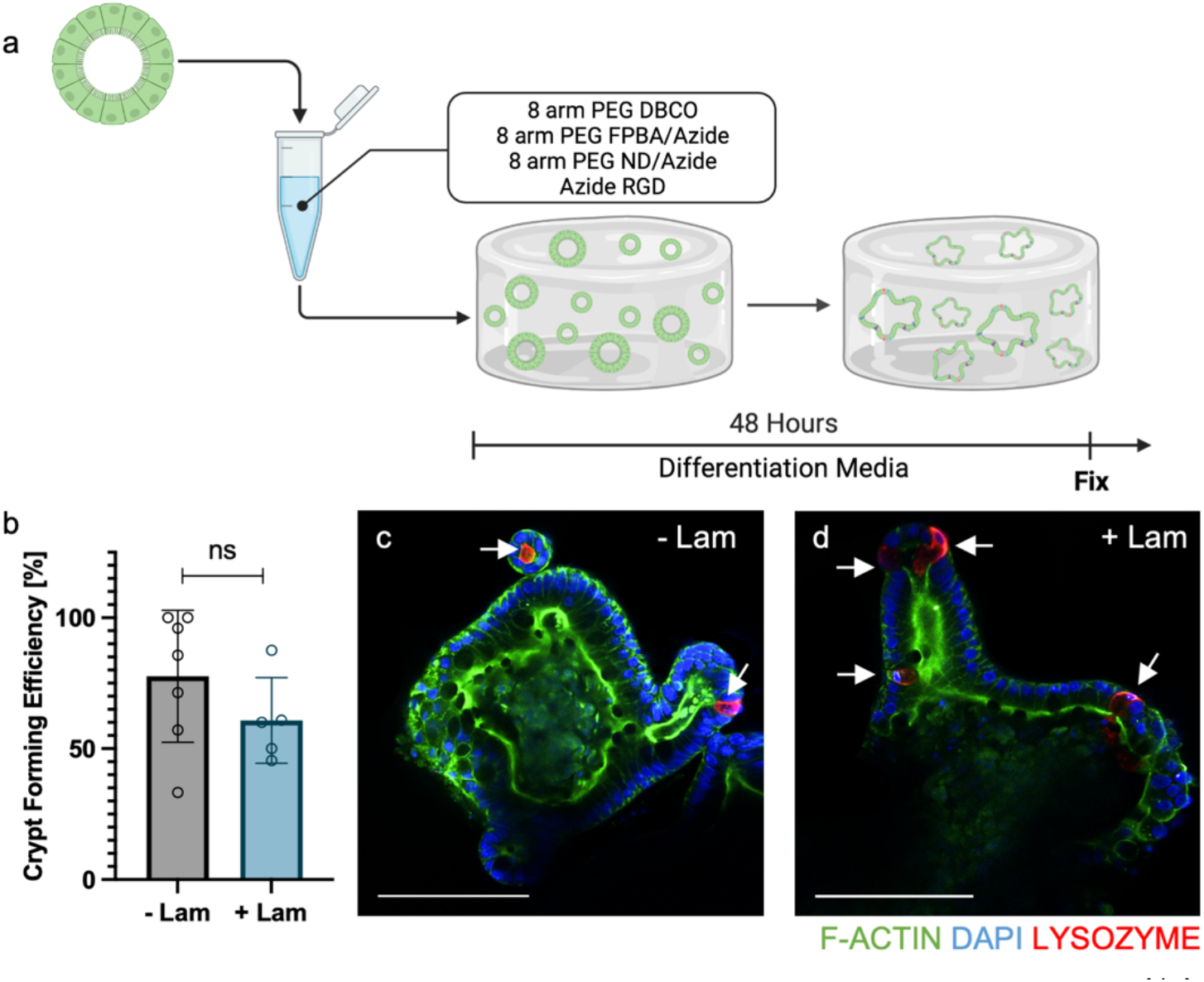
Boronate ester hydrogels promote intestinal organoid differentiation in the absence of supplemental laminin. **a**) Spherical, intestinal stem cell colonies were isolated from Matrigel and added to a solution containing 8-arm PEG dibenzocyclooctyne (DBCO) and azide functionalized RGD. 8-arm PEG fluorophenylboronic acid (FPBA)/azide and 8-arm PEG nitrodopamine (ND)/azide were simultaneously added, initiating spontaneously polymerization of the hydrogel network. The samples were cultured in differentiation media and after 48 hours the intestinal organoids were fixed. **b)** The crypt forming efficiency of intestinal organoids cultured in boronate ester hydrogels without exogenous laminin was as robust (-Lam) as when they were cultured in hydrogels with added laminin (+Lam) (+ Lam (*n* = 7), - Lam (*n* = 5); mean ± s.d., *ns* ≥ 0.05, one-way ANOVA). **c)** Intestinal organoid cultured in hydrogels without supplemental laminin (-Lam) and **d)** with supplemental laminin (+Lam). Scale bar 100 μm, white arrows denote location of Paneth cells in crypts.

### Laminin-organoid interactions influence crypt formation and architecture

To understand the role of organoid-laminin interactions in intestinal organoid differentiation, we encapsulated organoids in boronate ester hydrogels with and without exogenously added laminin and cultured them in differentiation media containing function-blocking antibodies against the α6 or β4 integrin subunits to block organoid-cell interactions with laminin (**Fig. 3a**).^16^ It is worthwhile to note that if laminin is not added to the matrix, the cells comprising the organoid have the capacity to secrete and interact with nascently deposited laminin.^10^ The α6β4 integrin is both relevant and specific as the β4 integrin subunit is integral in crypt fission^17^, a process similar to crypt formation in which one crypt splits into two, and the α6β4 integrin exclusively binds laminin.^18-24^ In the absence of supplemental laminin and presence of function-blocking antibodies, the CFE significantly decreased from 75% to 25% (α6) and 18% (β4) (**Fig. 3b)**. In the presence of exogenously added laminin and function-blocking antibodies, the CFE similarly decreased from 60% to 15% (α6) and 15% (β4). Collectively, these results implicate a crucial role for laminin in organoid crypt formation. We next sought to investigate the effect of laminin interactions on crypt architecture. When laminin is exogenously added to the matrix and laminin interactions are blocked with α6β4 integrin antibodies, the few crypts that form are of shorter length (i.e., ∼95 μm vs. ∼40 μm) and with fewer crypt per colony (from an average of 3 to 0.5) **(Fig. 3c,d)**. Similar changes are observed when laminin is not supplemented and α6β4 integrins are blocked.

**Fig 3.**
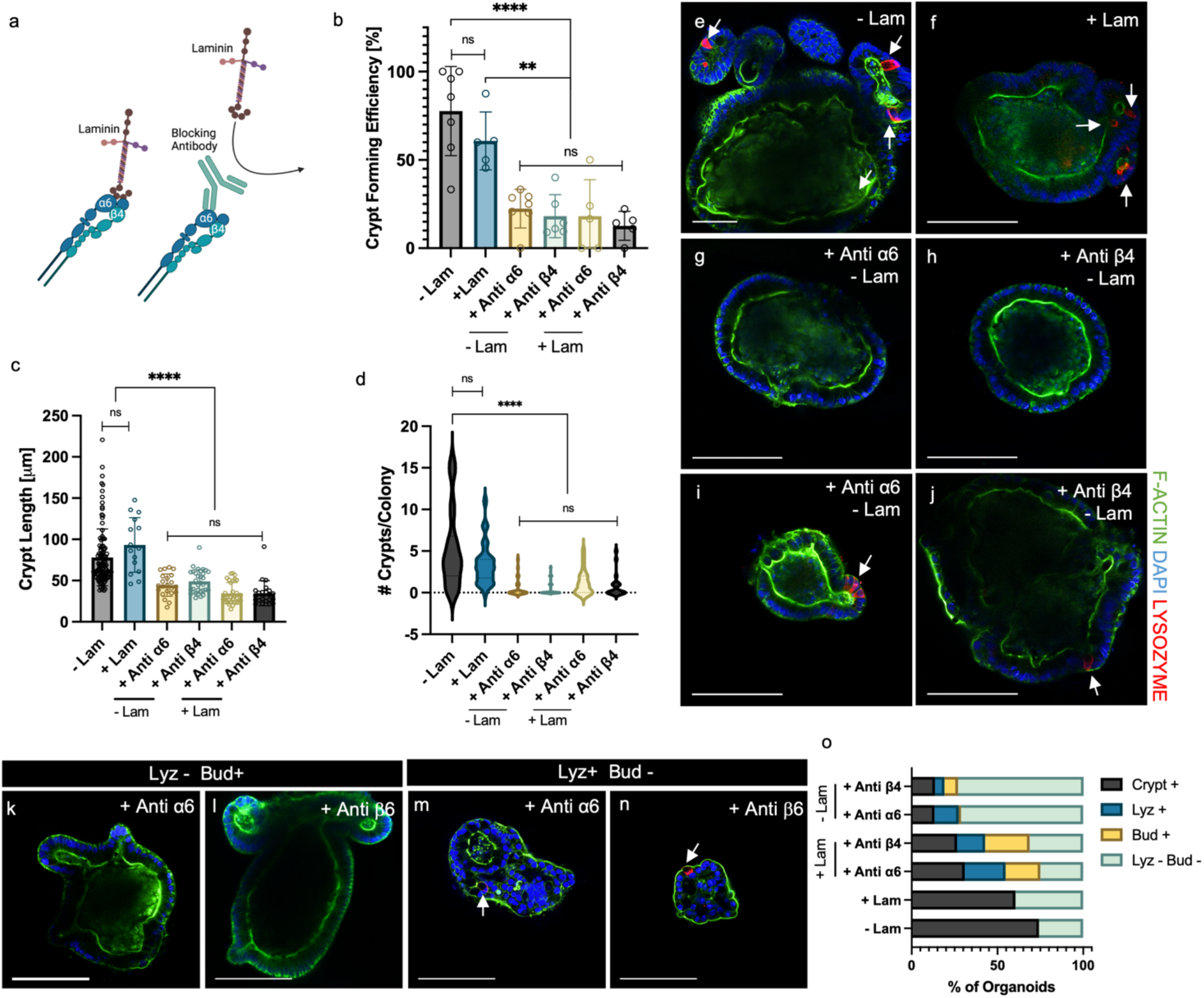
Laminin-organoid interactions influence measures of crypt architecture. **a)** Intestinal organoids largely interact with laminin via the α6β4 integrin. When either an α6 integrin subunit blocking antibody (+ Anti α6) or a β4 integrin subunit blocking antibody (+Anti β4) are added to media, organoid interactions with laminin are inhibited **b)** and the crypt forming efficiency of organoids in boronate ester hydrogels dramatically decreases, whether exogenous laminin has been added to the matrix (+ Lam) or not (-Lam) (– Lam (*n* = 7 hydrogels) and + Lam (*n* = 5 hydrogels) and are from Fig 2. +Anti α6 – Lam (*n =* 7 hydrogels), +Anti β4 – Lam (*n* = 6 hydrogels), +Anti α6 + Lam (*n* = 5 hydrogels), + Anti β4 + Lam (*n* = 5 hydrogels), mean ± s.d. *ns* ≥ 0.05, *\*\*P* ≤ 0.01, *\*\*\*\*P* ≤ 0.0001, one-way ANOVA) **c)** For organoids with blocked laminin interactions that still formed crypts, the crypts were smaller (*n* = 110 crypts analyzed for – Lam, 15 for + Lam, 22 for + Anti α6 – Lam, 31 for + Anti β4 – Lam, 31 for + Anti α6 + Lam, and 22 for + Anti β4 + Lam, each from at least 3 hydrogels per condition, mean ± s.d., *ns* ≥ 0.05, *\*\*\*\*P* ≤ 0.0001, one-way ANOVA) and **d)** there were fewer crypts per colony (*n* = 32 organoids analyzed for – Lam, 30 for + Lam, 28 for + Anti α6 – Lam, 34 for + Anti β4 – Lam, 21 for + Anti α6 + Lam, and 10 for + Anti β4 + Lam, each from at least 3 different hydrogels, violin plot, *ns* ≥ 0.05, *\*\*\*\*P* ≤ 0.0001, one-way ANOVA). **e)** The majority of colonies cultured in boronate ester hydrogels without (-Lam) or **f)** with (+Lam) formed budded intestinal organoids with Paneth cells (red) residing in their crypts. **g)** The majority of colonies grown in boronate ester hydrogels with anti α6 antibodies or **h)** anti β4 blocking antibodies remained cystic with no Paneth cells. A much smaller population of colonies encapsulated in hydrogels **i)** with anti α6 antibodies or **j)** with anti β4 antibodies formed fewer and smaller crypts compared to conditions without blocking antibodies. **k,l)** However, some organoids that were treated with blocking antibodies formed buds without Paneth cells (Lyz – Bud +) or **m,n)** had Paneth cells without budding (Lyz + Bud -) Scale bar 100 μm, white arrows point to Paneth cells in crypts. **o)** However, these populations were not seen in conditions without blocking antibodies. In this analysis, we define a crypt as a budding feature with at least one Paneth cell present. Scale bar 100 μm, white arrows point to Paneth cells in crypts (– Lam (*n =* 6 hydrogels), + Lam (*n* = 5 hydrogels),+ Anti α6 + Lam (*n* = 2 hydrogels), + Anti β4 + Lam (*n* = 3 hydrogels), + Anti α6 – Lam (*n* = 3 hydrogels), and + Anti β4 – Lam (*n* = 2 hydrogels), data presented as means).

In both - Lam conditions (**Fig. 3e**) and + Lam conditions (**Fig. 3f)**, organoids cultured in boronate ester hydrogels primarily contained crypts, defined as a feature composed of a bud with at least one Paneth cell. When interactions were blocked with either integrin subunit α6 or β4 blocking antibodies, the majority of organoids were cystic colonies that lacked budding and Paneth cells (**Fig. 3g,h**, all -Lam). The few organoids that did form crypts in these conditions had small, low density budding with Paneth cells **(Fig. 3i,j)**. Interestingly, there were also populations in the organoids with laminin interactions blocked that budded without Paneth cells present (**Fig. 3k,l**). Or, there were Paneth cells present without budding (**Fig. 3m,n**). These two unique populations were not present in the conditions with laminin interactions, implicating laminin’s role in the successful orchestration of symmetry breaking events in crypt formation (**Fig. 3o**). All in all, these studies demonstrate not only that laminin-organoid interactions influence crypt morphometrics (e.g., length and number of crypts), but are also required to successfully form crypts.

To validate laminin’s influence on these downstream aspects of crypt formation, we treated intestinal organoids with an inhibitor of focal adhesion kinase (FAK), a component of focal adhesions.^25-28^ In intestinal organoids, focal adhesions transduce ECM interactions to the nucleus to influence cell responses, including differentiation.^7^ Intestinal organoids were encapsulated (with or without exogenous laminin, + Lam and – Lam, respectively) in boronate ester hydrogels and cultured in media containing a FAK-14 inhibitor^26^ (**Fig. 4a-d**). We found the inhibition of FAK resulted in a decrease in crypt forming efficiency (**Fig. 4e**) and a reduction in the crypt size and number (**Fig. f,g**), similar to when laminin interactions were blocked via α6β4 blocking antibodies. The similarly impaired crypt formation when organoids were treated with FAK or blocked from interacting with laminin strengthens the connection between organoid and laminin interactions via focal adhesions and displays the influence of this signaling pathway on intestinal organoid crypt formation.

**Fig 4.**
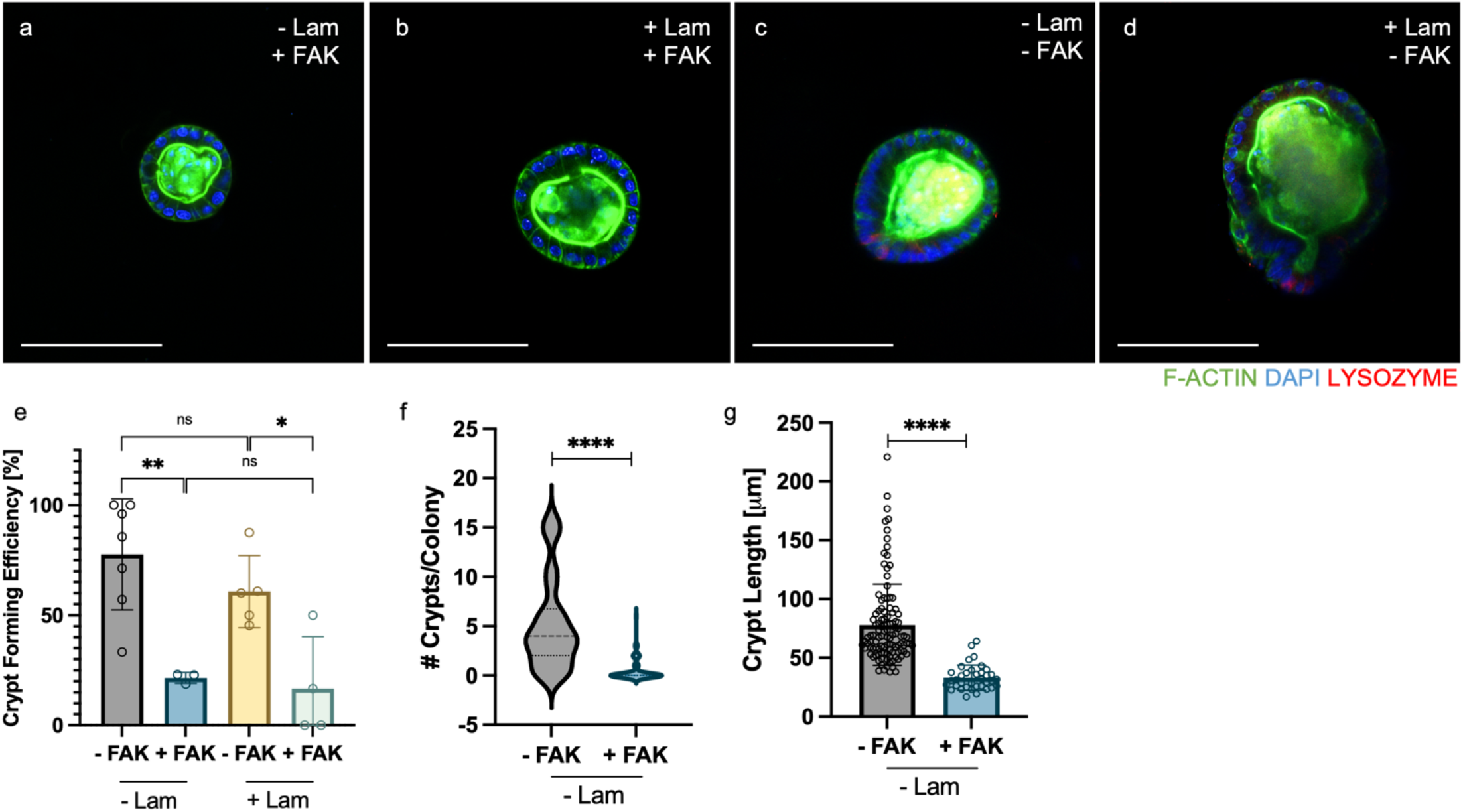
FAK inhibitor decreases crypt forming efficiency and influences crypt architecture. **a-d)** Intestinal organoids were encapsulated in boronate ester hydrogels with or without supplemental laminin (+/-Lam) and with or without the FAK inhibitor (+/-FAK) Scale bar 100 μm. Organoids were immunostained for the presence of crypts, marked by Paneth cells (red). **e)** When intestinal organoids were cultured in boronate ester hydrogels with a FAK inhibitor, crypt forming efficiency decreased regardless of whether laminin was added to the matrix (+ Lam) or not (-Lam) (*n* = 7 hydrogels for – FAK – Lam, 3 for + FAK – LAM, 5 for – FAK + LAM and 4 for + FAK + Lam, - FAK condition data from Fig 3, mean ± s.d., *ns* ≥ 0.05, *\*P* ≤ 0.05, *\*\*P* ≤ 0.01, one-way ANOVA). **f)** Similarly, the number of crypts per colony decreased (*n* = 32 organoids analyzed for – FAK – Lam and 77 for + FAK - Lam, both from 3 hydrogels per condition, - FAK condition data from Figure 3, violin plot, *\*\*\*\*P* ≤ 0.0001, one-way ANOVA) and **g)** the size of colonies decreased (*n* = 110 crypts analyzed for – FAK – Lam and 36 for + FAK - Lam, both from 3 hydrogels per condition, - FAK condition data from Figure 3, mean ± s.d., *\*\*\*\*P* ≤ 0.0001, one-way ANOVA).

### Intestinal organoids deposit laminin in boronate ester hydrogels

Given the α6β4 integrin solely binds laminin, our function blocking antibody data suggest that the organoids in the boronate ester hydrogels without supplemental laminin must be depositing their own laminin to elicit these ECM interactions guiding crypt formation. To investigate this hypothesis, we encapsulated intestinal organoids in boronate ester hydrogels without supplemental laminin and immunostained with a pan specific laminin antibody. We discovered that intestinal organoids deposited robust laminin matrix (**Fig. 5a**) with a thicker laminin matrix in crypt regions compared to the organoid body (**Fig. 5b**). Additionally, more of the crypt surface was covered in laminin compared to the body, with percent coverage at 75% and 15%, respectively (**Fig. 5c**). These results are notable as the crypt is the source of intestinal stem cells in intestinal organoids and the location of the initial stages of organoid differentiation. There was no laminin matrix present at the time of encapsulation. (**Supplemental Fig. 1**).

**Fig 5.**
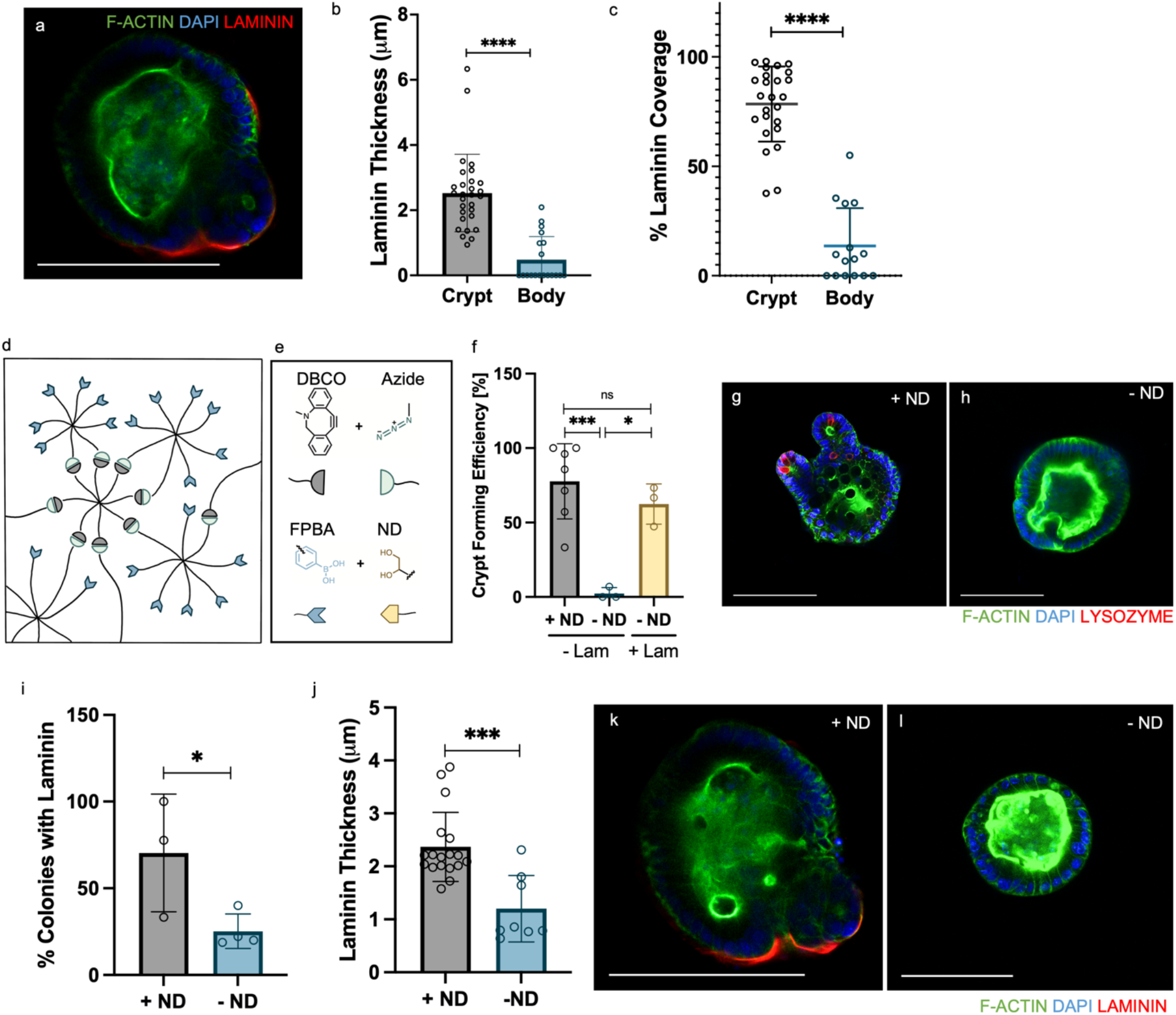
Intestinal organoids deposit laminin in boronate ester hydrogels. **a)** Intestinal organoids cultured in boronate ester hydrogels deposited laminin (scale bar 100 μm), **b)** with more laminin in crypt regions compared to the organoid body *(n* = 29 organoids analyzed for crypt and 19 for body, both from 3 hydrogels per condition, mean ± s.d., *\*\*\*\*P* ≤ 0.0001, unpaired t-test). **c)** The percent of crypt surface that was covered by laminin was similarly greater than the percent of organoid body that was covered by laminin (*n* = 25 organoids analyzed for crypt and 15 for body, both from 3 hydrogels per condition, mean ± s.d., *\*\*\*\*P* ≤ 0.0001, unpaired t-test). **d)** A variation in the boronate ester hydrogel formulation that has only permanent bonds and no adaptable bonds was implemented for organoid culture. This system does not contain the 8-arm PEG nitrodopamine (ND)/azide and only contains the 8-arm PEG DBCO and 8-arm PEG fluorophenylboronic acid (FPBA) /azide arms. **e)** In this system, the DBCO and azide groups form covalent bonds via the SPAAC reaction (gray and green, respectively), and because there is no ND present (yellow), the FPBA is free and the matrix is not stress relaxing, but elastic. **f)** Organoids were encapsulated in the elastic (-ND) and stress relaxing (+ND) forms of the boronate ester hydrogels, with (+Lam) or without (-Lam) supplemental laminin. Organoids grown in the elastic hydrogel (-ND) had decreased crypt forming efficiency when laminin was not present, but the crypt formation was rescued when laminin was added to the matrix (+ ND – Lam (*n* = 7 hydrogels), – ND – Lam (*n* = 3 hydrogels), - ND + Lam (*n* = 3 hydrogels), +ND condition data from Fig 2, mean ± s.d., *\*P* ≤ 0.05, *\*\*\*P* ≤ 0.001, one-way ANOVA). **g,h)** Organoids were immunostained for lysozyme, DAPI and f-actin to evaluate for the presence of crypts as marked by Paneth cells (red) in +ND and –ND hydrogels (both without Lam). Scale bars 100 μm. **i)** The percentage of organoid colonies with associated laminin was lower in the elastic (-ND) compared to the stress relaxing (+ND) hydrogels, (+ ND (*n* = 3 hydrogels), - ND (*n* = 4 hydrogels), mean ± s.d., *\*P* ≤ 0.05, unpaired t-test) and **j)** The thickness of the secreted laminin near organoid crypts was thinner in elastic (-ND) compared to the stress relaxing (+ND) hydrogels (*n* = 29 organoids for + ND and 19 for – ND both from 3 hydrogels per condition, mean ± s.d. *\*\*\*P* ≤ 0.001, unpaired t-test). **k,l)** These laminin interactions were evaluated through the immunostaining of intestinal organoids for Laminin (Red), DAPI and F-actin in stress relaxing (+ ND) and elastic (-ND) hydrogels. Scale bars 100 μm.

To enable differentiation in the absence of supplemental laminin in boronate ester hydrogels, we hypothesized that the network’s rapid stress relaxation may increase laminin deposition, as it has long been established that viscoelasticity influences aspects of cell-matrix interactions, including matrix deposition.^7,8,13,14,29-34^ To that end, we implemented an elastic variation of the boronate ester hydrogels to be compared to the viscoelastic version. In this system, the 8-arm PEG ND/azide is omitted (**Fig. 5d**), and the remaining two macromers (8-arm PEG DBCO, 8-arm PEG FPBA/azide) can only form a covalent, SPAAC network, via reaction of the DBCO and azide groups (**Fig. 5e)**. In the absence of ND, the FPBA groups are free and do not interact with any other groups, thus eliminating the stress relaxing nature of the original network.

When intestinal organoids were encapsulated in these elastic boronate ester hydrogels (-ND), we find that the organoids do not form crypts in the absence of supplemental laminin (**Fig. 5f-h)**, but when laminin is added that CFE is rescued (**Fig. 5f**). Further, there was significantly less laminin deposited in the elastic hydrogels compared to the stress relaxing condition (+ND) (**Fig. 5i)**, with 25% of colonies containing deposited laminin compared to almost 75% of organoids in the viscoelastic hydrogels. Similarly, the colonies that did deposit laminin in the elastic hydrogels had a thinner laminin matrix than the viscoelastic condition (**Fig. 5j**-**l**). Therefore, we demonstrate a dynamics dependent deposition of laminin matrix in intestinal organoids. While FPBA can interact with diol groups, which are present on ECM proteins like laminin, the excess, free FPBA groups in the elastic hydrogel were not sufficient for crypt formation without supplemented laminin. Therefore, we can conclude that sequestering of cell-secreted laminin via interactions with FPBA does not explain the ability of intestinal organoids to form crypts without exogenous laminin in boronate ester hydrogels. Further, if laminin was being sequestered, we would expect laminin to be less mobile in the elastic hydrogel, but it is equally immobile in both elastic and viscoelastic hydrogels (**Supplemental Fig. 2)**. Taken together, our data suggest that the unique adaptable bond dynamics and viscoelasticity of the boronate ester hydrogels increases laminin deposition to the extent that laminin does not need to be added exogenously for crypt formation.

### Viscoelasticity influences nascent protein deposition in intestinal organoids

To investigate if the influence of viscoelasticity on the secretory profile of organoids translates to proteins beyond laminin, we employed a technique for fluorescent labeling of non-canonical amino acids.^35^ Intestinal organoids were encapsulated in stress relaxing boronate ester hydrogels with methionine free differentiation media supplemented with an alkyne containing methionine analog, L-homoproparglyglycine (HPG). Newly deposited proteins were labeled with a fluorescent azide through a copper catalyzed azide alkyne cycloaddition reaction (**Fig. 6a**). Organoids were additionally treated with a plasma membrane stain to visualize cell borders (**Fig. 6b**) to ensure only extracellular protein was being analyzed (**Fig. 6c**).^35^ Analogous to our observed laminin dynamics, we discover when cultured in viscoelastic boronate ester hydrogels, organoid crypt regions contained thicker deposited matrix than the organoid body (**Fig. 6d**). Similarly, the surface coverage of the crypt in deposited ECM was higher than in the body, with over 75% of the crypt covered in ECM compared to under 50% of the organoid body (**Fig. 6e**).

**Fig 6.**
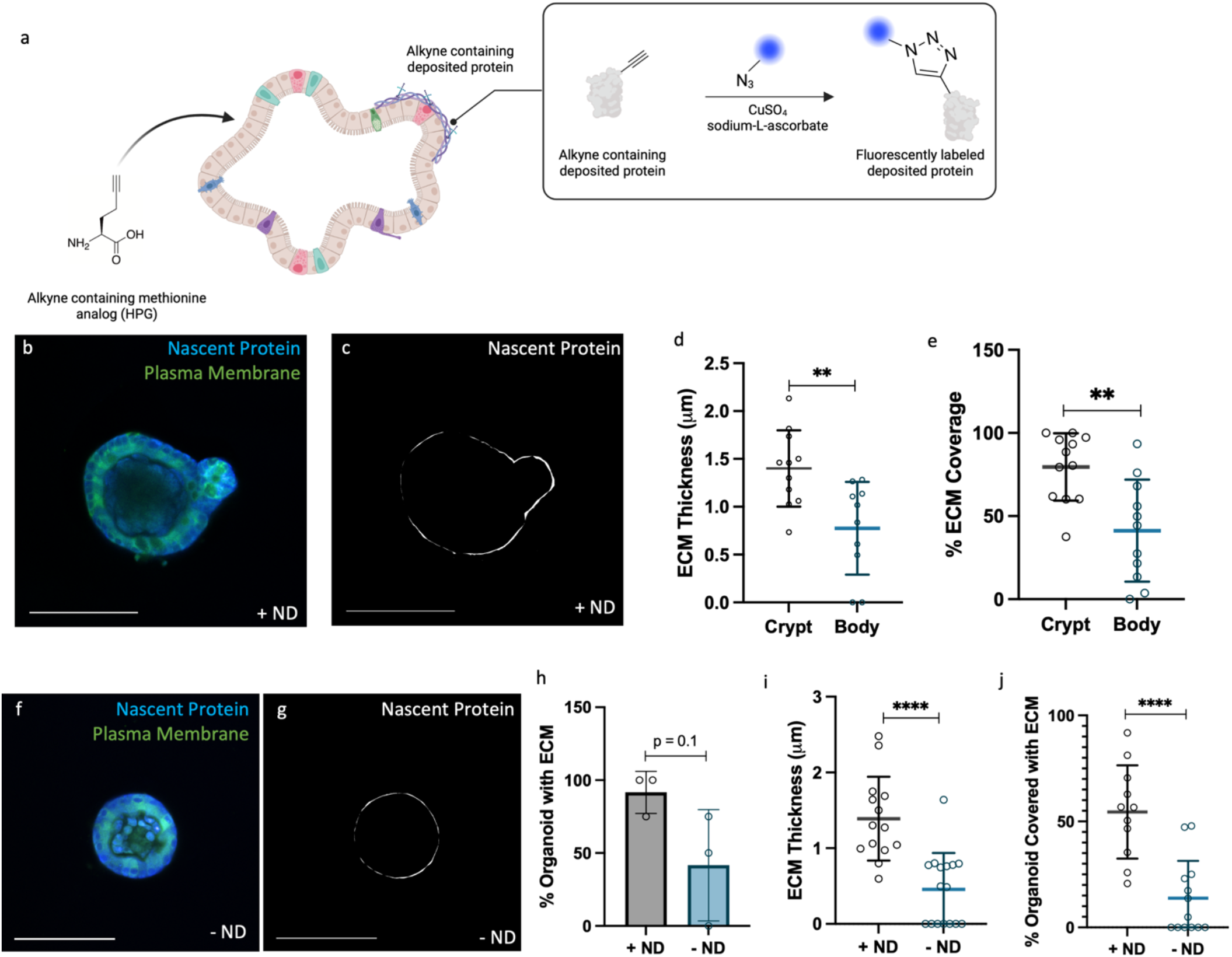
Stress relaxation allows for increased intestinal organoid nascent protein deposition. **a)** Intestinal organoids were cultured in boronate ester hydrogels in methionine free base media that was supplemented with an alkyne containing methionine analog. The organoid uptakes this analog and secretes proteins containing alkyne groups. These alkyne groups can be fluorescently labeled with an azide fluorophore through a copper catalyzed SPAAC reaction. **b)** Deposited protein from intestinal organoids cultured in stress relaxing boronate ester hydrogels (+ND) was labeled with a 405 azide fluorophore (blue) and stained with CellMask Plasma Membrane Green, **c)** to ensure that only extracellular protein was analyzed (white) Scale bar 100 □m. **d)** Organoids had more deposited protein in the crypt region compared to the body of the organoid (*n* = 11 organoids analyzed from 3 independent biological experiments, mean ± s.d., *\*\*P* ≤ 0.01, unpaired t-test). **e)** The percent of crypt surface covered with deposited protein was greater in the crypt regions (*n* = 12 organoids analyzed for crypt and 11 for body, both from 3 independent biological experiments, mean ± s.d., *\*\*P* ≤ 0.01, unpaired t-test). **f)** The elastic (-ND) boronate ester hydrogel formulation without the 8-arm PEG nitrodopamine/azide was again implemented to evaluate the influence bond adaptability on nascent protein deposition. Nascent protein was fluorescently labeled blue and organoids were stained with CellMask Plasma Membrane Green **g)** to allow for visualization of extracellular protein (white) Scale bars 100 □m. **h)** A lower percentage of organoids in the elastic (-ND) hydrogels deposited matrix than those in stress relaxing (+ND) hydrogels (*n* = 3 independent biological experiments, mean ± s.d., unpaired t-test), and **i)** the organoids in the stress relaxing (+ND) hydrogels deposited a thicker matrix (*n* = 14 organoids analyzed for + ND and 13 for - ND, both from 3 independent biological experiments, mean ± s.d., *\*\*\*\*P* ≤ 0.0001, unpaired t-test). **j)** Similarly, the percentage of the organoid surface covered in ECM was greater in the stress relaxing (+ND) compared to the elastic (-ND) hydrogel *(n* = 11 organoids analyzed for + ND and 13 for - ND, both from 3 independent biological experiments, mean ± s.d., *\*\*\*\*P* ≤ 0.0001, unpaired t-test).

To compare this deposition to that of intestinal organoids in elastic hydrogel environments, we again encapsulated intestinal organoids in hydrogels that were void of the 8-arm PEG ND/azide macromer. Fewer organoids in the elastic (- ND) system deposited nascent protein compared to those in the viscoelastic (+ND) system (**Fig. 6f-h**). Intestinal organoids grown in viscoelastic hydrogels deposited thicker protein matrices and had more of their surface covered in ECM, with 60% coverage compared to ∼30% in elastic hydrogels (**Fig. j**). This secretory dependence on viscoelasticity helps elucidate why synthetic hydrogels to date have required supplementation of laminin, as all networks historically utilized for organoid culture have been elastic or had longer stress relaxation time scales and lower extents of relaxation then the boronate ester hydrogels. All together these studies establish the influence of viscoelasticity on nascent protein deposition and interaction in intestinal organoids and establish their key role in intestinal organoid morphogenesis.

## OUTLOOK

Intestinal organoids hold vast potential for clinical applications such as screening drugs, personalizing medicine and transplanting tissue. Two major bottlenecks for realizing organoid clinical potential are (1) the biological components (i.e., Matrigel, laminin) comprising the materials used to culture them and (2) the gaps in understanding how ECM signaling influences organoid morphogenesis. The former increases the potential for immune reactions for transplantation and decreases the control over the material properties and consequently the organoid structure and function. The latter limits the ability to engineer improved materials with precise user control that facilitate predictable and controlled growth of organoids, a requirement for the stringent regulations of clinical applications.

In this work, we successfully culture intestinal organoids in a fully synthetic hydrogel system without exogenous biological factors like Matrigel and laminin, lending inspiration to future design of fully synthetic materials for intestinal organoid growth. Fully synthetic hydrogel systems inherently allow greater user control of the cellular environment than their biological counterparts and will impart greater predictability and versatility for organoid studies and clinical applications. We further use the boronate ester system to establish dynamics (e.g., adaptable bonds, stress relaxation) dependent secretion of nascent protein and demonstrate organoid interactions with these spatially varying proteins influence crypt formation and structure. An understanding of organoid-hydrogel and the consequent organoid-protein interactions is similarly useful in designing controlled presentation of differentiation cues to intestinal organoids.

Intriguingly, other stress relaxing matrices employed to culture intestinal organoids required exogenous laminin addition for intestinal organoid differentiation. *Chaudhuri et al*. argue that the effect of viscoelasticity on cell behavior is linked to the concept of mechanical confinement.^32,33^ When cellular processes such as motility, proliferation, migration or matrix deposition are limited by physical restrictions in three dimensions, the cells are considered mechanically confined. In elastic and rigid hydrogels, this confinement is overcome by matrix degradation strategies. In contrast, increasing the rate of stress relaxation allows cells to increasingly overcome confinement to deposit matrix proteins, proliferate and migrate. Therefore, we posit that the uniquely rapid relaxation of the boronate ester matrix is a defining difference that allows for nascent protein deposition to promote organoid differentiation and crypt formation without exogenous laminin. All together, these observations implore future investigation into the influence of viscoelasticity on intestinal organoid development and differentiation. As we learn more about the mechanical and biological factors that influence organoid development, we can engineer improved and precisely defined matrices to efficiently grow functional organoids, facilitating their translation and exciting potential applications.

## METHODS

### Synthesis of hydrogel macromers

8-arm PEG dibenzocylcooctyne (20 kDa), 8-arm PEG fluorophenylboronic acid/azide (20 kDa) and 8-arm PEG nitrodopamine/azide (20 kDa) were synthesized as previously described.^13^ Our syntheses yielded 64% functionalized 8-arm PEG dibenzocylcooctyne, a 8-arm PEG nitrodopamine/azide product with 5 arms functionalized with nitrodopamine and 3 with azide, and a 8-arm PEG fluorophenylboronic acid/azide product containing 3 arms functionalized with fluorophenyl boronic acid and 5 arms with azide. Functionalization was determined with HNMR and TNBSA assays.

### Characterization of hydrogels

Stock solutions of 10 wt% 8-arm PEG DBCO (20 kDa), 20 wt% 8-arm PEG fluorophenylboronic acid/Azide (20 kDa) and 20 wt% 8-arm PEG nitrodopamine/azide were prepared using phosphate buffered saline (PBS). Hydrogels were formed on stoichiometry by simultaneously adding 8-arm PEG fluorophenylboronic acid/azide and 8-arm PEG nitrodopamine/azide to an open syringe barrel that already contained 8-arm PEG DBCO. The solution was titrated and stirred and then allowed to polymerize for 15 minutes before being ejected into a PBS bath, where it was swollen to equilibrium for 30 min. Hydrogels were transferred to the bottom plate of a shear oscillatory rheometer (DHR-3, TA Instruments) fitted with an 8 mm parallel plate geometry. The geometry was lowered until an axial force of 0.2 N was reached, at which point mineral oil was applied to surround the gel. Storage modulus was recorded at 5% strain and 1 Hz. To quantify stress relaxation, the hydrogels were brought to a 10 % strain and the stress was recorded.

### Crypt isolation and organoid culture

Murine small intestinal crypts were isolated from Lgr5-eGFP-IRES-CreERT2 mice as previously described.^3^ The resulting crypts were cultured as organoids in reduced growth factor Matrigel (Corning). Organoids were kept as stem colonies in Advanced Dulbecco’s Modified Eagles Medium (DMEM)/F-12 (Invitrogen) containing N2 and B27 supplements (Thermo Fisher Scientific), GlutaMax (Gibco), Hepes and penicillin-streptomycin supplemented with epidermal growth factor (EGF, “E,” 50 ng/mL, R&D Systems), Noggin (“N,” 100 ng/mL, PeproTech), R-spondin-conditioned medium (“R,” 5% v/v, Organoid & Tissue Modeling Shared Resource, CU Anschutz), CHIR99021 (“C,” 3 μM, Sell-eckchem), and Valproic Acid (“V,” 1 mM, Sigma-Aldrich). Media containing Advanced DMEM/F-12, N2, B27, Hepes and penicillin-streptomycin is called “basal medium.” Organoids were passaged every 4 days and media was changed every 2 days. The University of Colorado Institutional Animal Care and Use Committee approved the animal protocol (no. 00084) for this research.

### Encapsulation of colonies into boronate ester hydrogels

Two days prior to encapsulation, stem colonies were passaged into Matrigel with aggressive shearing prior to encapsulation in Matrigel. Over the two days, the colonies would reform into cystic, stem colonies. On the day of encapsulation, colonies were released from Matrigel using ice-cold DMEM/F-12 and collected in a conical tube. The conical tube was centrifuged for 4 min at 900 rpm to form a pellet of colonies. The DMEM/F-12 was aspirated and replaced with a small volume of fresh DMEM/F-12 to resuspend the colonies. The cell solution was added to a diluted volume of 8-arm PEG DBCO and azide functionalized Arg-Gly-Aps (RGD 0.8 mM) in an open syringe barrel. 8-arm PEG fluorophenylboronic acid/azide and 8-arm PEG nitrodopamine/azide were added simultaneously, titrated and stirred. All precursor solutions were kept on ice until used. The syringes were transferred to a 37 ºC incubator and allowed to polymerize for 15 minutes. The hydrogels (30 μL in size) were ejected into wells of a 48 well plate filled with 400 μL of basal medium with EGF, R-Spondin and Noggin at the concentrations listed in the previous section (ENR) to promote differentiation. Organoids were cultured for 48 hours and then fixed and immunostained.

### Fixation and immunofluorescence staining of intestinal organoids

Media was removed from wells containing organoid laden boronate ester hydrogels and replaced with 4% paraformaldehyde for 30 min at room temperature on a rocker. Organoids were then permeabilized with 0.2 % Triton X-100 (Sigma Aldrich) for 1 hour. The samples were then incubated overnight at 4 ºC in blocking buffer containing 10% Goat Serum (Thermo Fisher) and 0.01% Triton X-100. Primary antibodies were added at 4 ºC overnight (Rabbit anti Lysozyme, Thermo Fisher Scientific PA129680, 1:50; Rabbit anti laminin, Abcam ab11575, 1:100). Samples were washed 5 x 1 hours with PBS at 4 ºC. Then secondary antibodies were then added in blocking buffer over night at 4 ºC (Goat anti Rabbit 647, Thermo Fisher Scientific A21245, 1:1000; Rhodamine Phalloidin, Fisher Scientific 50-646-256, 1:200; DAPI, Sigma Aldrich 10236276001, 1:2000 of 2 mg/mL solution). Samples were then washed at 4 x 1 hour with PBS at room temperature and stored at 4 ºC in PBS while awaiting imaging.

### Function blocking antibody treatment

Intestinal organoids were encapsulated into boronate ester hydrogels and placed into differentiation media containing 10 μg/mL of blocking antibody (anti integrin α6 antibody BD Pharmingen #555734 or anti integrin β4 antibody BD Pharmingen #553745). At 48 hours samples were fixed and immunostained as described above.

### FAK inhibitor 14 treatment

A 3 mM solution of Fak Inhibitor 14 (Millipore Sigma SML0837) was made using PBS. Intestinal organoids were encapsulated in boronate ester hydrogels as described above and placed in basal medium + ENR with FAK Inhibitor 14 added to a final concentration of 10 μM. After 48 hours the sampled were fixed and immunostained as described above.

### Fluorescent labeling of non-canonical amino acids

Methods for fluorescent labeling of non-canonical amino acids were adapted from *Loebel et al*. 2022.^35^ Custom methionine free DMEM/F-12 media was ordered from Thermo Fisher and prepared with GlutaMax (Gibco), Hepes, penicillin-streptomycin, and N2 and B27 supplements (Thermo Fisher Scientific). Media was supplemented with a mixture of L-homoproparglyglycine (HPG, Click chemistry tools 354232) and methionine (Millipore Sigma M5308) (total methionine content of 75% and 25%, respectively). Intestinal organoids were switched to using this media (including ENRCV, same concentrations as above) at least one passage prior to encapsulation. Organoids were released from Matrigel and encapsulated into boronate ester hydrogels as previously described. Organoids were cultured in the methionine analog supplemented media with ENR for 48 hours. Media was removed and samples were treated with 4% PFA for 30 min and washed 3 x 10 min with PBS. Samples were treated with 1 μL per 400 μL of PBS of a monofunctionalized PEG-azide for 30 min on a rocker to cap any unreacted DBCO groups on the 8-arm PEG DBCO in the boronate ester network. A solution of 2% bovine serum albumin was prepared with PBS and used to wash samples 3 x 5 minutes on a rocker at room temperature. CellMask Plasma Membrane Stain Green (Thermo Fisher Scientific C37608) was diluted 1:500 in 2% BSA and added to the samples for 2 hours at room temperature on a rocker. The CellMask was removed and samples were washed 3 x 5 minutes with 2 % BSA. For the copper catalyzed fluorescent labeling of HPG, a solution of Tris buffered saline with 19.8 mg/mL sodium-L-ascorbate (Millipore Sigma 11140) and a solution of 400 mM CuSO_4_ in Tris buffered saline were prepared. The CuSO_4_ solution was added to the sodium-L-ascorbate solution to a final concentration of 4 mM. To this solution, azide fluorophore (AZDye 405 Azide, Click Chemistry Tools 1307-5) to a concentration of 30 μM. Samples were treated with the copper catalyzed fluorescent labeling solution overnight at 4 ºC. Samples were washed heavily, at least 4 x 1 hour with PBS and then imaged or stored at 4 ºC while awaiting imaging.

### Imaging and image analysis

Samples were imaged using confocal microscopy (Zeiss LSM 710) and analysis was carried out using FIJI. The line measuring tool was used to measure crypt length. Thickness of laminin was analyzed by separating the F-actin and laminin fluorescent channels in FIJI and thresholding the image slices. The F-actin channel was used to create a mask that was subtracted from the laminin channel to ensure only extracellular laminin was analyzed. The Ridge Detection Plug-In was then used to quantify average laminin thickness as a function of location around the organoid circumference. Similarly, to calculate the coverage of organoids in ECM, the organoids were additionally stained with a CellMask Plasma Membrane stain and imaged on a confocal microscope. The channel slices were split and thresholded, and the CellMask stain was used to make a mask that was subtracted from the nascent protein channel to eliminate any protein in the organoid. The thickness as a function of location around the organoid circumference was analyzed using the Ridge Detection Plug-In in FIJI.

### Statistics

To assess statistical significance, Prism 9 (GraphPad) was used to conduct unpaired student *t* tests for experiments with two conditions and one-way ANOVA were used with more than two conditions. *p* < 0.5 were considered significant. All experiment replicates were conducted as noted in the text.

## Acknowledgements

Schematics in figures were created using BioRender.com. This work was supported by a grant from the National Institutes of Health R01 DK120921 (K.S.A.), P30-DK116073 (P.J.D.), P30-CA046934 (P.J.D.), K99 DK135907-01 (M.R.B.) and F31 DK126427 (F.M.Y.); DARPA grant W911NF-19-2-004 (L.J.M., B.E.K.); and National Science Foundation RECODE 2033723 (K.S.A.) and National Science Foundation Graduate Research Fellowship Program DGE 2040434 (E.A.H.).

