## Supplementary Information for "Fully synthetic hydrogels promote robust crypt formation in intestinal organoids"

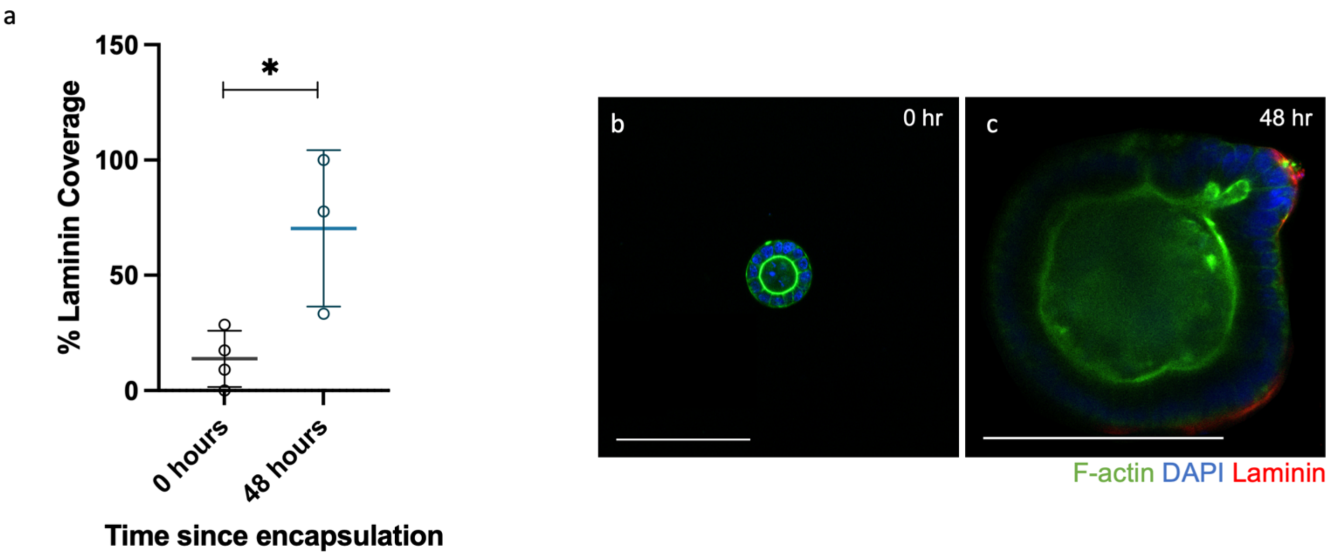


**Supplemental Fig 1. Intestinal organoids do not have a laminin matrix when encapsulated in boronate ester hydrogels. a)** The percent of organoid surface covered with laminin at the time of encapsulation is minimal compared to the amount of laminin present 48 hours post encapsulation once the organoids have deposited their own matrix. ($n$ = 4 hydrogels for 0 hours and 3 for 48 hours, mean ± s.d., *$P \leq 0.05$, unpaired t-test.) **b)** At zero hours the intestinal organoids encapsulated were cystic colonies with no laminin present compared to **c)** the organoids 48 hours after encapsulation when robust laminin matrix had been deposited (red, scale bar 100 μm).

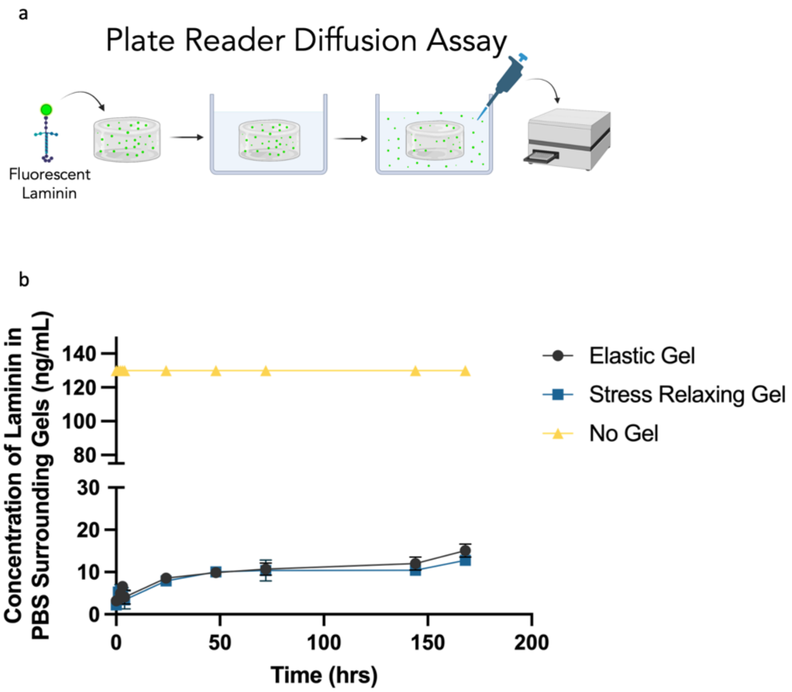


**Supplementary Fig 2. Laminin does not diffuse out of the boronate ester hydrogels. a)** Fluorescent laminin was encapsulated boronate ester hydrogels where the “stress relaxing gel” is the stress relaxing boronate ester hydrogel and the “elastic gel” is an 8-arm PEG DBCO and 8-arm PEG azide. The hydrogels were put in wells containing PBS and samples were taken from the PBS baths at regular time intervals. The fluorescence of those samples were measured on a plate reader and used to calculate the concentration of laminin. **b)** Concentration of laminin in the PBS bath was measured as a function of time, and very minimal laminin diffused out of the elastic and stress relaxing hydrogels. The “no gel” condition was a bath of PBS with laminin at the same concentration as laminin was encapsulated in the hydrogels to control for any decrease in laminin fluorescence with time.
